# Intercellular mitochondrial transfer and transmitophagy in response to protein import dysfunction

**DOI:** 10.1101/2025.11.10.687620

**Authors:** Emily Glover, Beth Wiseman, Celyn Dugdale, Lorna Hodgson, Kevin Wilkinson, Ian Collinson

## Abstract

Mitochondrial protein import is critical for organelle biogenesis and maintenance, essential processes for cellular homeostasis. Import dysfunction compromises cellular energy supplies which is damaging to cells, particularly those with high energetic demands like neurons. Previously, we have shown that import failure is rescued by intercellular mitochondrial transfer *via* tunnelling nanotubes (TNTs) however, the fate of the transferred mitochondria and the mechanistic basis for rescue was unresolved. Here, we show that bi-directional mitochondrial trafficking between cells harbouring import-defective and import-competent mitochondria has distinct regulation and consequences. Transferred import-defective mitochondria are highly fragmented and destined for canonical lysosomal degradation. In contrast, ROS-producing mitochondria at the periphery of cells with import-competent mitochondria are transferred into neighbouring cells undergoing import failure; these new arrivals then accumulate within previously uncharacterised ‘mitochondrial degradation bodies’ (MBDs). We speculate that the co-operation of these distinct examples of TNT-mediated conventional and non-canonical ‘transmitophagy’ instigate mitochondrial regeneration, and thereby rescue.

## Introduction

Intercellular mitochondrial transfer can have a significant and sustained beneficial impact on recipient cell metabolism (Baldwin *et al*, 2024; Islam *et al*, 2012; Lin *et al*, 2024; Hoover *et al*, 2025; Marlein *et al*, 2017, 2019; Brestoff *et al*, 2021), even leading to rescue of cells containing defective mitochondria (Spees *et al*, 2006; Rostami *et al*, 2017; Needs *et al*, 2024). However, in most previous studies it is unclear how transferred mitochondria have been able to elicit these effects.

Exogenous mitochondria can have a diversity of fates, including incorporation into (Crewe *et al*, 2021), or the replacement of (Ikeda *et al*, 2025), the endogenous network, in addition to the segregation from the endogenous network as a functional mitochondrial sub-population (Baldwin *et al*, 2024; Masuzawa *et al*, 2013). Surprisingly though, most studies show that exogenous mitochondria remain segregated from the endogenous network and display a small and spherical morphology (Lin *et al*, 2024; Islam *et al*, 2012; Kidwell *et al*, 2023; Vallabhaneni *et al*, 2012; Wang & Gerdes, 2015; Yao *et al*, 2018), sometimes clustered in the perinuclear region (Lin *et al*, 2024; Islam *et al*, 2012). Notably, these are characteristic of lysosomal mitochondria (Frank *et al*, 2012; Pu *et al*, 2016), suggesting that these mitochondria are destined for degradation.

Recent reports suppose that exogenous mitochondria may function as instigators of signalling pathways in the receiving cell (Lin *et al*, 2024; Kidwell *et al*, 2023; Ikeda *et al*, 2025), explaining in part how small numbers of dysfunctional and short-lived exogenous mitochondria can be amplified to induce profound cell-wide responses. Other reports describe a different narrative, whereby intercellular mitochondrial transfer for degradation by a neighbouring cell—‘transmitophagy’ (Davis *et al*, 2014; Morales *et al*, 2020; Nicolás-Ávila *et al*, 2020)—whilst important for mitochondrial quality control of the donor cell, reaps no benefit the recipient (Nicolás-Ávila *et al*, 2020). It is therefore currently unclear how transcellular degradation of mitochondria can result in two distinctive outcomes – signalling or quality control – and how these functions might collaborate to mediate the metabolic rescue of recipient cells seen after intercellular mitochondrial transfer.

Previously, we have shown that cells subject to chronic mitochondrial import failure upregulate TNT-mediated intercellular mitochondrial transfer and this rescues the import defect (Needs *et al*, 2024). However, the fate of exogenous mitochondria post-transfer and how this instigates rescue was not determined. In this study, we demonstrate bi-directional transfer of import-defective and import-competent mitochondria between co-cultured cells and use flow cytometry to isolate cells containing exogenous mitochondria after this two-way traffic. Imaging by fluorescence and electron microscopy show that transferred import-defective mitochondria are exported for conventional mitophagy while transferred import-competent mitochondria are sequestered into a membrane-bound body for non-canonical degradation. Our findings suggest transmitophagy has both mitochondrial quality control and signalling roles, which together rescue cells harbouring import-defective mitochondria.

## Results & Discussion

### TNT-mediated intercellular mitochondrial transfer in response to mitochondrial import failure is bidirectional

Previously, we have found that the numbers of TNTs increased in response to chronic mitochondrial import inhibition conditions and that these can mediate intercellular mitochondrial transfer (Needs *et al*, 2024, 2023). However, the directionality of TNT-mediated mitochondrial transfer between cells containing import-defective mitochondria and cells containing import-competent mitochondria and their respective consequences were not fully explored.

To address these questions, HeLa cells were conditioned in galactose (HeLaGAL), rendering cells OXPHOS-dependent and thus sensitive to mitochondrial perturbation (Aguer *et al*, 2011; Marroquin *et al*, 2007). HeLaGAL cell lines were then created with stable expression of a precursor construct comprising the mitochondrial targeting sequence of subunit 9 of the ATP synthase from the fungus *Neurospora crassa* (su9) (Westermann & Neupert, 2000; Hartl *et al*, 1989), fused to the fluorescent proteins EGFP, mScarlet, or mTagBFP2 for visualisation. Dihydrofolate reductase (DHFR) was fused to the C-terminus of this precursor to create lines susceptible to mitochondrial import inhibition by virtue of DHFR resisting unfolding when bound to the small molecule methotrexate (MTX). Given that import requires precursor unfolding, this blocks the import machinery (Rassow *et al*, 1989; Eilers & Schatz, 1986; Needs *et al*, 2024) (Figure 1A).

**Figure 1:**
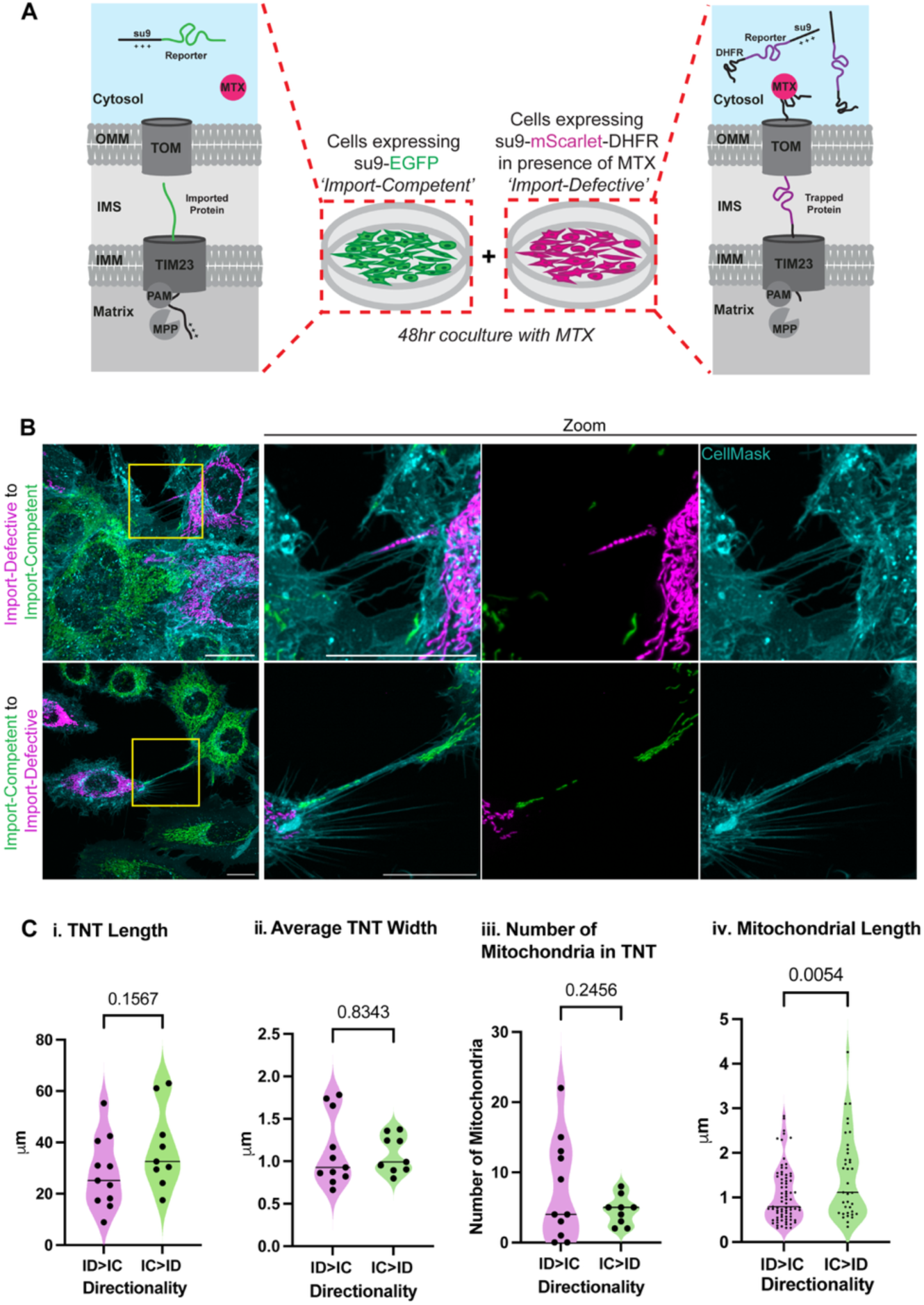
TNT-mediated intercellular mitochondrial transfer in response to mitochondrial import failure is bidirectional. (A) Schematic of coculture system. (B – top panel) Representative image of su9-mScarlet-DHFR (+ MTX) expressing mitochondria (import-defective, magenta) in a TNT. (B – bottom panel) Representative image of su9-EGFP expressing mitochondria (import-competent, green) in a TNT. All scale bars 20µm. (C) Quantification of 20 TNTs over N=3 biological repeats of; (i) TNT length, (ii) average TNT width, (iii) number of mitochondria in TNT, (iv) mitochondrial length. ID; import-defective, IC; import-competent. P values from unpaired T test.

To investigate the TNT-mediated transfer of import-defective *versus* import-competent mitochondria, su9-mScarlet-DHFR and su9-EGFP lines were co-cultured with MTX for 48hrs (Figure 1A). Live florescence imaging of mitochondria-containing TNTs (mitoTNTs), identified as a membranous tube containing mitochondria connecting a su9-mScarlet-DHFR-expressing cell and a su9-EGFP-expressing cell, revealed similar numbers of mitoTNTs containing import-defective mitochondria (55%) and mitoTNTs containing import-competent mitochondria (45%) (Figure 1B). This indicates that both import-competent and import-defective mitochondria are mobilised. Notably, import-defective and import-competent mitochondria were never in the same TNT at the same time, suggesting that mitochondrial transfer within an individual TNT is uni-directional. Irrespective of the state of the transiting mitochondria, mitoTNT length and width were very similar (Figure 1Ci-ii).

With respect to the frequency of the mitochondrial traffic within each TNT, there tended to be more import-defective mitochondria within each individual (mean of 7.55) in comparison to import-competent mitochondria (mean of 4.56) (Figure 1Ciii). Transiting import-defective mitochondria were significantly shorter than those that were import-competent (1.05μm and 1.47μm, respectively) (Figure 1Civ), consistent with the former being more prone to fission (Needs *et al*, 2024). Taken together, this data suggests that intercellular transfer of import-competent and import-defective mitochondria both make significant contributions to the rescue of chronic import inhibition previously observed (Needs *et al*, 2024).

### Cells subject to intercellular mitochondrial transfer can be isolated by florescence activated cell sorting

Flow cytometry approaches can be used to quantify and isolate cells that have received intercellularly transferred cargo (Kidwell *et al*, 2023; Ikeda *et al*, 2025; Lin *et al*, 2024; Kolba *et al*, 2019; Sharma & Subramaniam, 2019; Kumar *et al*, 2017). Therefore, we developed a flow cytometry assay in which import-defective and import-competent mitochondria were distinguished by different fluorescently labeled mitochondria. Cells subject to intercellular mitochondrial transfer could thus be identified and isolated for analysis.

Following gating of alive singlet cells using parental HeLaGALs (Figure 2A), a four-way gating system was set up using su9-mScarlet-DHFR and su9-EGFP mono-cultures (Figure 2B). A co-culture of su9-mScarlet-DHFR and su9-EGFP stable cell lines with MTX was then subject to this gating strategy.

**Figure 2:**
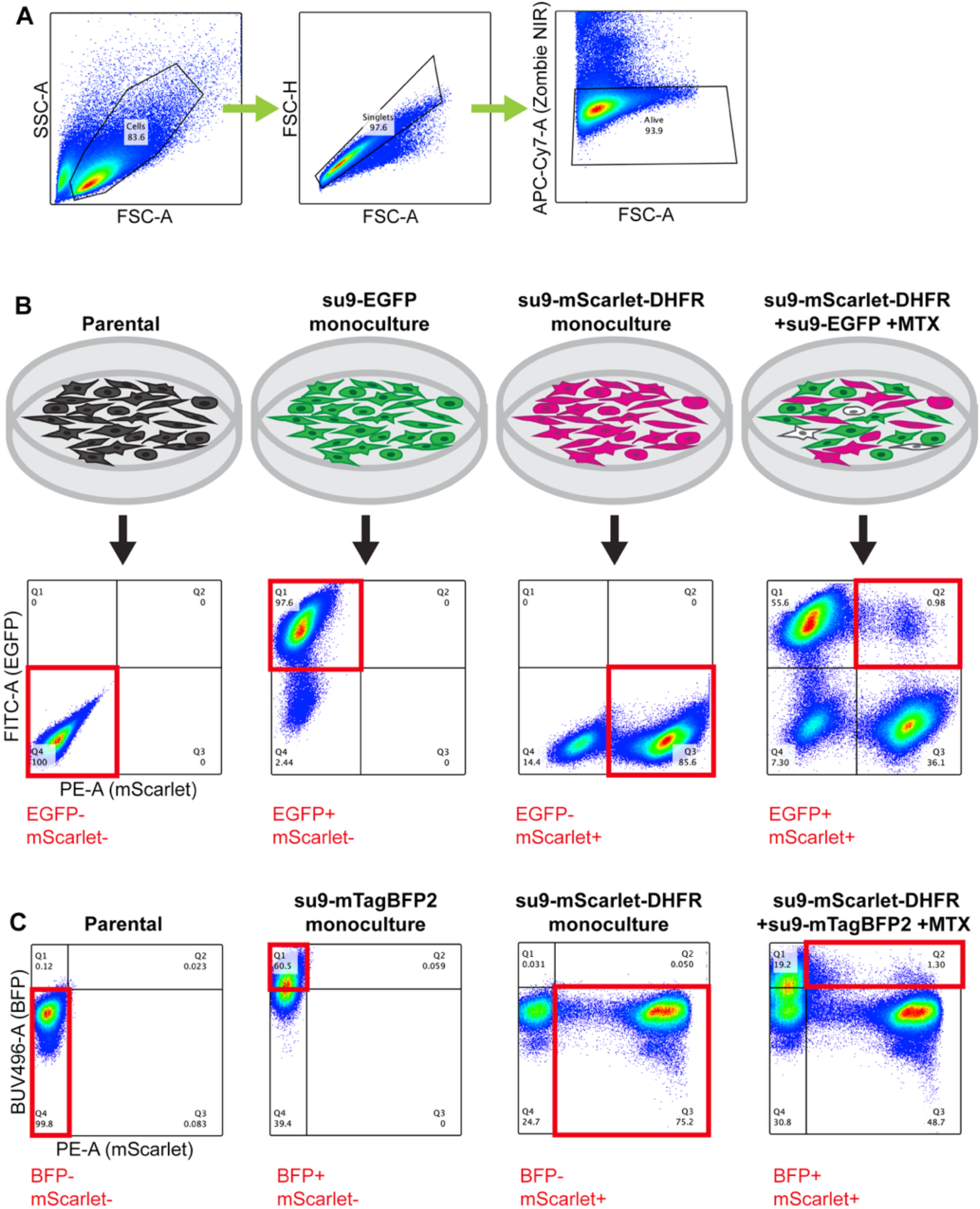
Cells subject to intercellular mitochondrial transfer can be isolated by florescence activated cell sorting. (A) Gating strategy for isolation of cells that are singlets and alive (B) four-way gating system for identification of EGFP+/mScarlet+ cells (C) four-way gating system for identification of BFP+/mScarlet+ cells.

The majority of live singlet cells in this co-culture were separated into either the EGFP only or mScarlet only gates: cells that have retained their resident mitochondria and had not undergone mixing. However, a population of cells were double positive, containing both EGFP and mScarlet, indicative of their containment of both import competent and compromised mitochondria—the result of mitochondrial transfer (Figure 2B).

Given the possibility of lysosomal encapsulation for degradation (mitophagy) of migrating mitochondria, and the resulting low pH, it became necessary to switch to a pH insensitive fluorophore in order to monitor their fate. Therefore, EGFP, which is quenched in the acidic environment of the lysosome was exchanged for mTagBFP2, which is not (Kim & Seong, 2021; Shinoda *et al*, 2018). Importantly, for the purposes of our analysis, mTagBFP2 is subject to lysosomal proteolysis (Shinoda *et al*, 2018), whilst mScarlet is neither quenched nor degraded (Clancy *et al*, 2023). Therefore, lysosomal degradation of mitochondria could be characterised by the gradual loss of mTagBFP2 fluorescence.

A four-way gating system analogous to that used above was set up using su9-mScarlet-DHFR and su9-mTagBFP2 mono-cultures (Figure 2C). Our analysis showed this was capable, albeit at a slight loss in resolution, of distinguishing cells containing both mTagBFP2 and mScarlet for sorting.

### Transferred import-competent and import-defective mitochondria have distinct fates

To investigate the consequences of mitochondrial transfer, a co-culture of su9-mScarlet-DHFR and su9-mTagBFP2 expressing cell lines were grown in the presence of MTX. Cells double positive for mScarlet and mTagBFP2 were then sorted and subject to live fluorescence imaging (Figure 3A).

**Figure 3:**
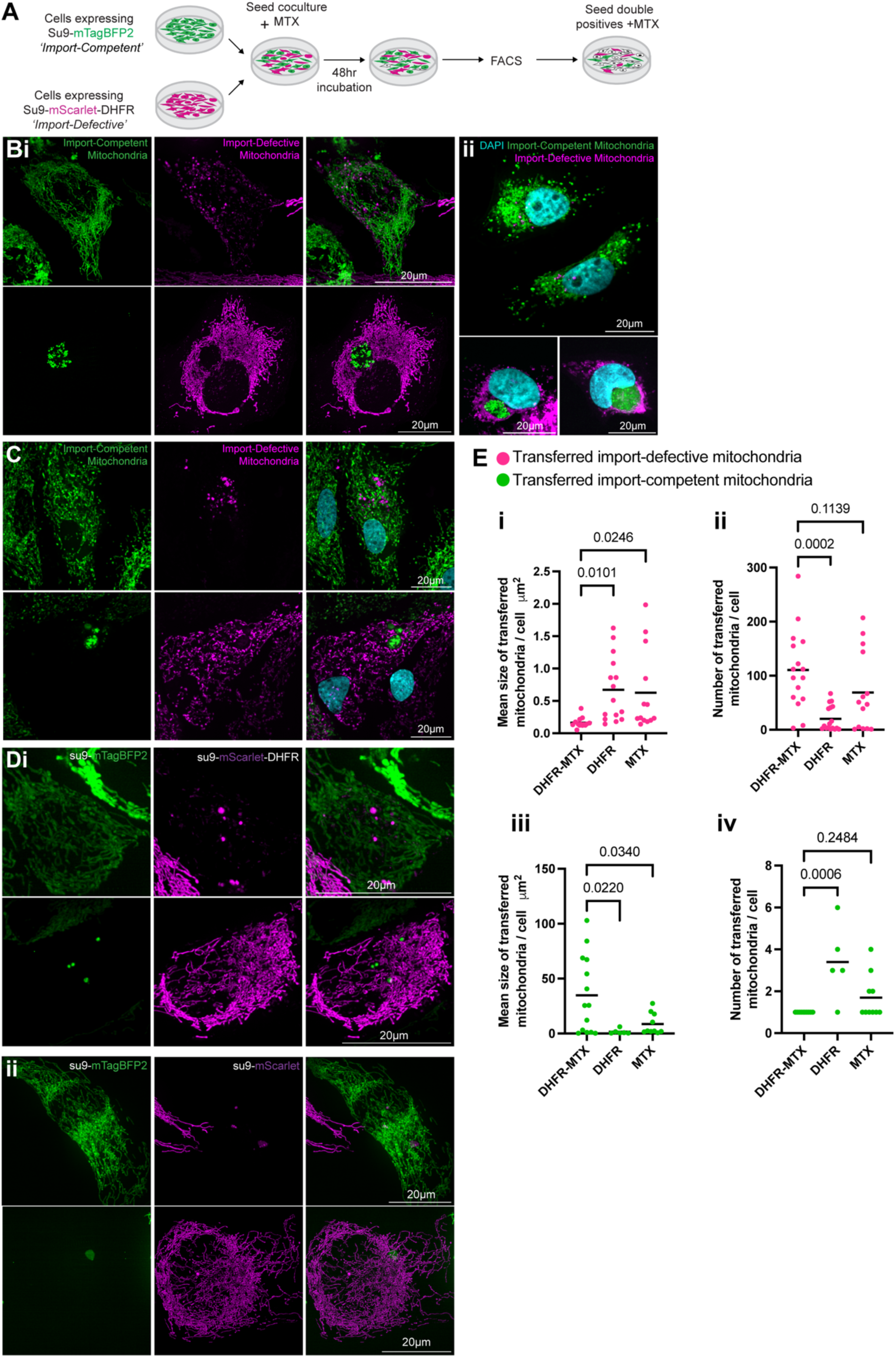
Transferred import-competent and import-defective mitochondria have distinct fates. (A) Schematic of workflow. (Bi) Representative images of double positive cells from a 48hr coculture of su9-mTagBFP2 and su9-mScarlet-DHFR cell lines in the presence of MTX. (Bii) Representative images of double positive cells from a 48hr coculture of su9-EGFP and su9-mScarlet-DHFR cell lines in the presence of MTX following fixation and DAPI staining, confirming cells containing transferred mitochondria are mono-nuclear. (C) examples of double positive cells from a coculture of primary astrocytes expressing either su9-EGFP-DHFR or cox8a-DsRed. (Di) Representative images of double positive cells from a 48hr coculture of su9-mTagBFP2 and su9-mScarlet-DHFR cell lines. (Dii) Representative images of double positive cells from a 48hr coculture of su9-mTagBFP2 and su9-mScarlet cell lines in the presence of MTX. (E) Quantification of mean size of transferred mitochondria and number of transferred mitochondria per cell, taken from max. projected images. For transferred import-competent mitochondria localised in mitochondrial degradation bodies (MDBs), individual mitochondria could not be distinguished so the total MDB area was quantified. n=62 cells from N=3 biological replicates. P values from unpaired T test.

Interestingly, two distinctive types of cells were observed (Figure 3Bi). In the first of these, cells containing import-competent mitochondria (su9-mTagBFP2) were the recipients of numerous (mean of 111) import-defective mitochondria (su9-mScarlet-DHFR) which were small (mean of 0.163 µm^2^), spherical and dispersed throughout the cell (Figure 3Bi, top panel). The second type was cells containing a network of import-defective mitochondria (su9-mScarlet-DHFR) with the addition of a defined cluster of import-competent mitochondria (su9-mTagBFP2) – referred to hereafter as a ‘mitochondrial degradation body’ MDB) (Figure 3B, bottom panel).

On average, a singular extraordinarily large MBD (mean of 34.8 µm^2^) could be seen in each import-defective cell. DAPI staining confirmed that these MDBs were devoid of chromatin and, therefore, unlikely to have arisen by cell-cell fusion, and consistent with them being the product of intercellular mitochondrial transfer (Figure 3Bii). Furthermore, we observed analogous structures in a co-culture of import-competent rat primary astrocytes (cox8a-DsRed) alongside those with import-defective mitochondria (su9-GFP-DHFR) (Figure 3C). Therefore, these distinctive MDBs are unlikely to be an artefact of the clonal cell lines examined thus far, but potentially a previously uncharacterised organelle.

Taken together, the results show that intercellular mitochondrial transfer occurs bidirectionally between cells populated by import-competent and import-defective mitochondria. Additionally, the transferred mitochondria do not fuse and become incorporated into the recipient mitochondrial network, consistent with what has been observed elsewhere (Lin *et al*, 2024; Ikeda *et al*, 2025; Islam *et al*, 2012; Kidwell *et al*, 2023; Vallabhaneni *et al*, 2012; Wang & Gerdes, 2015; Yao *et al*, 2018; Baldwin *et al*, 2024). Whilst the appearance of transferred import-defective mitochondria within the recipient is typical of the small and spherical morphology of transferred mitochondria seen previously (Kidwell *et al*, 2023; Vallabhaneni *et al*, 2012; Wang & Gerdes, 2015; Yao *et al*, 2018; Lin *et al*, 2024; Islam *et al*, 2012), the appearance of MDBs upon the counter-flow of import-competent mitochondria was entirely unexpected.

To verify that these effects were the consequences of mitochondrial import inhibition, the impacts of DHFR or MTX removal from the co-culture were assessed. Cells expressing su9-mTagBFP2 were co-cultured together with either *(i)* su9-mScarlet-DHFR in the absence of MTX (Figure 3Di) or *(ii)* su9-mScarlet with MTX (Figure 3Dii). Double positive cells were then isolated and compared to those from su9-mTagBFP2 and su9-mScarlet-DHFR co-cultures in the presence of MTX.

With regards to transferred mScarlet labelled mitochondria, analysis showed that their small size (mean of 0.163 µm^2^) was dependent on blocked import sites (presence of DHFR and MTX), since DHFR or MTX alone resulted in a larger size (mean of 0.671µm^2^ and 0.626µm^2^, respectively) (Figure 3Ei). Moreover, import-defective mitochondria have a higher frequency of mitochondrial transfer: on average 111/cell, compared to DHFR (20/cell) or MTX (∼69/cell) utilised alone (Figure 3Eii).

With respect to transferred import-competent mitochondria (visualised by mTagBFP2), their accumulation into large MDBs, characterised by low numbers and very large size, was dependent on their entry into cells subject to mitochondrial import site blocking (presence of DHFR and MTX) (Figure 3Eiii-iv). When the recipients were exposed to a DHFR containing precursor or MTX separately, the transferred import-competent mitochondria were less inclined to be segregated into MDBs, as demonstrated by their larger number and smaller size (Figure 3Eiii-iv).

Therefore, intercellular mitochondrial transport still occurs in co-cultures wherein both cell lines contain import-competent mitochondria (DHFR or MTX omitted). However, the transfer of a large number of highly fragmented mitochondria as well as the trafficking of mitochondria into MDBs are specific features of bidirectional transfer between cells with import-competent mitochondria and cells undergoing mitochondrial import failure.

### Transferred import-defective and import-competent mitochondria undergo distinct forms of transmitophagy

In order to collect more detailed information on the fate of transferred mitochondria, we used correlative light and electron microscopy (CLEM). This showed that transferred import-defective mitochondria have a classic lysosomal morphology, indicative of conventional transmitophagy (Figure 4Aii-iii, red arrows). In contrast, MDBs resulting from transfer of import-competent mitochondria into cells undergoing mitochondrial import failure, are packed with membranous material (Figure 4Bii-iii). Double membrane-bound structures can be seen within (Figure 4Bii, green arrows), which are presumably mitochondria that undergoing lysosomal degradation (Figure 4Bii, blue arrows). Furthermore, a membrane can be seen at the edge of the MDBs (Figure 4 Biii, yellow arrows), which could be part of an encapsulating structure.

**Figure 4:**
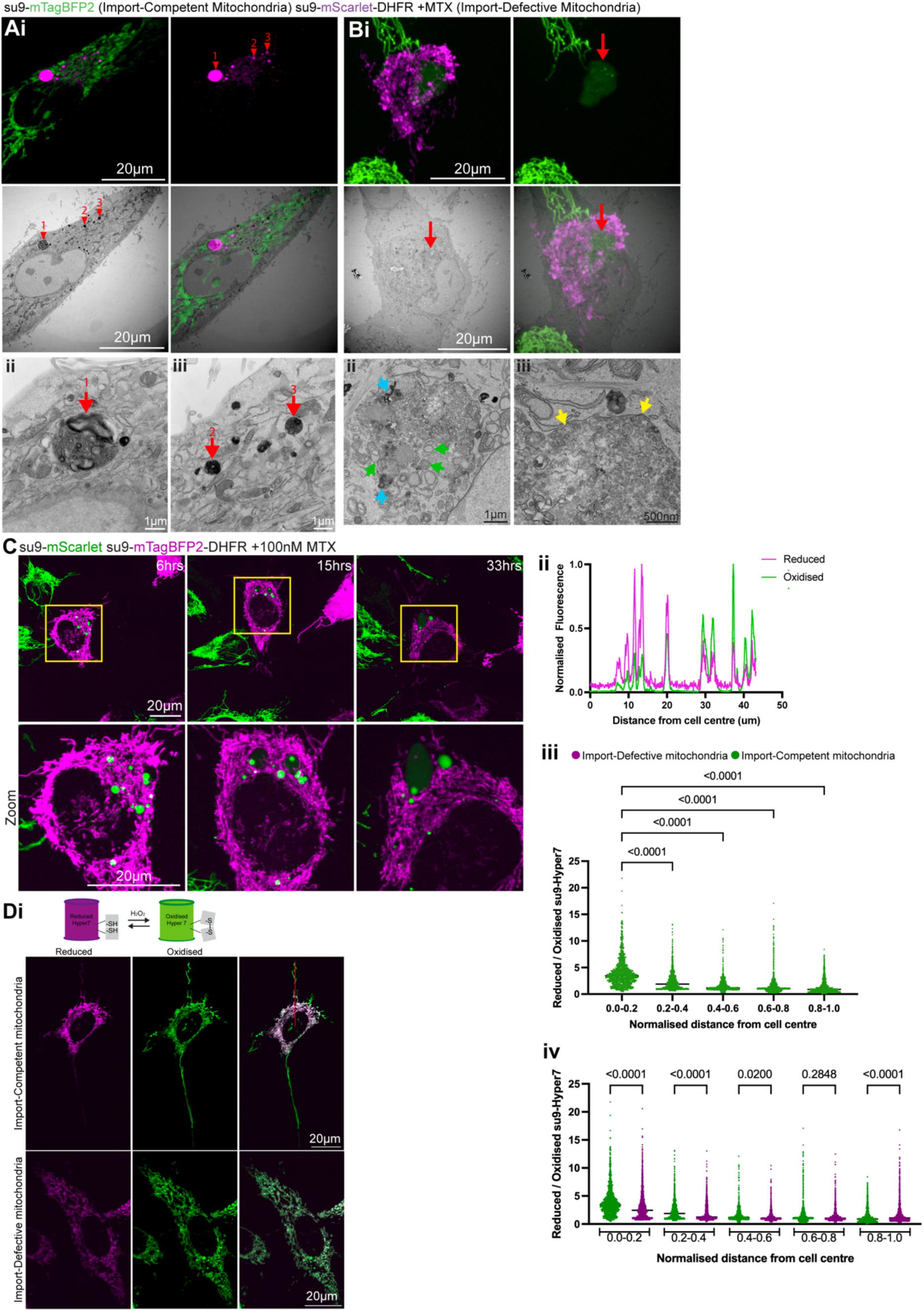
Transferred import-defective and import-competent mitochondria undergo distinct forms of transmitophagy. (Ai – top panel) Florescence microscopy image of transferred import-defective mitochondria (red arrows), (Ai– bottom panel) correlated with electron microscopy image. (ii and iii) Electron microscopy of transferred lysosomal import-defective mitochondria. (Bi – top panel) Florescence microscopy image of mitochondrial degradation body (MDB) (red arrow), (Bi – bottom panel) correlated with electron microscopy image. (Bii) electron microscopy of MDB containing potential ex-mitochondria (green arrows), lysosomes (blue arrows) and (Biii) surrounded by a partial membrane (yellow arrows). (C) Stills from a movie of transferred import-competent mitochondria (su9-mScarlet) in a cell containing import-defective mitochondria (su9-mTagBFP2-DHFR +100nM MTX) at indicated time points post-sorting, demonstrating MDB formation (Di) Schematic of Hyper7 oxidation by hydrogen peroxide and representative image of su9-Hyper7 HeLaGAL line (top panel, import-competent mitochondria) and su9-Hyper7;su9-mScarlet-DHFR cell line in the presence of 100nM MTX (bottom panel, import-defective mitochondria). (Dii) normalized fluorescence intensities of emission band pass (BP) 525/50 in response to excitation at 405nm (reduced Hyper7) or excitation at 488nm (oxidized Hyper7) vs. distance from cell centre (Diii) reduced/oxidized su9-Hyper7 ratio in HeLaGAL cell line vs. normalised distance from cell centre. n=17 cells over N=3 biological repeats. (Div) reduced/oxidized su9-Hyper7 ratio in HeLaGAL cell line (import-competent mitochondria) in comparison to HeLaGAL cell line transfected with su9-mScarlet-DHFR and treated with 100nM MTX for 48hrs (import-defective mitochondria) vs. normalised distance from cell centre. n=17-19 cells over N=3 biological repeats. P values from one-way ANOVA.

In order to gain time resolved information regarding mitochondrial incorporation into MDBs, we performed live imaging on double positive cells isolated from a su9-mScarlet and su9-mTagBFP2-DHFR cocultured in the presence of MTX. This showed that transferred import-competent mitochondria were initially present with a fragmented appearance analogous to those of import-defective mitochondria that have moved in the other direction (Figure 4C, 6hr time point). Then, over time they cluster into increasingly large MDBs (Figure 4C, 33hr time point) in a process presumably instigated by the receiving cell undergoing mitochondrial import failure.

These observations pose the question of why transferred import-competent mitochondria are trafficked into MDBs, while transferred import-defective mitochondria remain dispersed. To address this, we analysed matrix ROS in HeLaGAL cells by the production of mitochondrially-targeted ROS sensor HyPer7 (Pak *et al*, 2020). Interestingly, we observed that peripheral mitochondria in import-competent HeLaGAL cells have a significantly more oxidised mitochondrial matrix relative to those in the perinuclear region (representative image in Figure 4Di, top panel and quantification in ii and iii). This distribution places oxidised mitochondria in prime position for intercellular transfer. Notably, Kidwell e*t al.* show that ROS production by transferred mitochondria can activate ERK signalling in the receiving cell (Kidwell *et al*, 2023). Therefore, we propose that ROS production by mitochondria transferred from the periphery of import-competent cells into those undergoing mitochondrial import failure promotes their trafficking into MBDs for non-canonical mitophagy in the receiving cell.

In contrast, while perinuclear mitochondria in HeLaGAL cells harbouring import-defective mitochondria display a more oxidised matrix, the peripheral mitochondria in these cells are more reduced compared to HeLaGAL cells containing non-compromised organelles (representative image in Figure 4Di, bottom panel and quantification in iv). We suggest that the export of these mitochondria with relatively lower ROS production leads to more conventional lysosomal mediated mitophagy, rather than by the formation of MDBs.

### Transferred mitochondria are ultimately degraded

To understand the long-term fate of transferring mitochondria, extended live imaging of cells double was performed on double positive cells isolated from a coculture of import-competent and import-defective mitochondria. As noted above, both mScarlet and mTagBFP2 remain visible in the lysosome, but only the latter can be degraded (Kim & Seong, 2021; Shinoda *et al*, 2018; Clancy *et al*, 2023). Thus, when cells expressing su9-mTagBFP2-DHFR were co-cultured with those expressing su9-mScarlet, we were able to follow the fate of transferred import-defective mitochondria—by virtue of their fluorescence loss (Figure 5Ai). Conversely, a co-culture of su9-mScarlet-DHFR and su9-mTagBFP2 cell lines was used to follow the fate of transferred import-competent mitochondria within the MDBs (Figure 5Aii).

**Figure 5:**
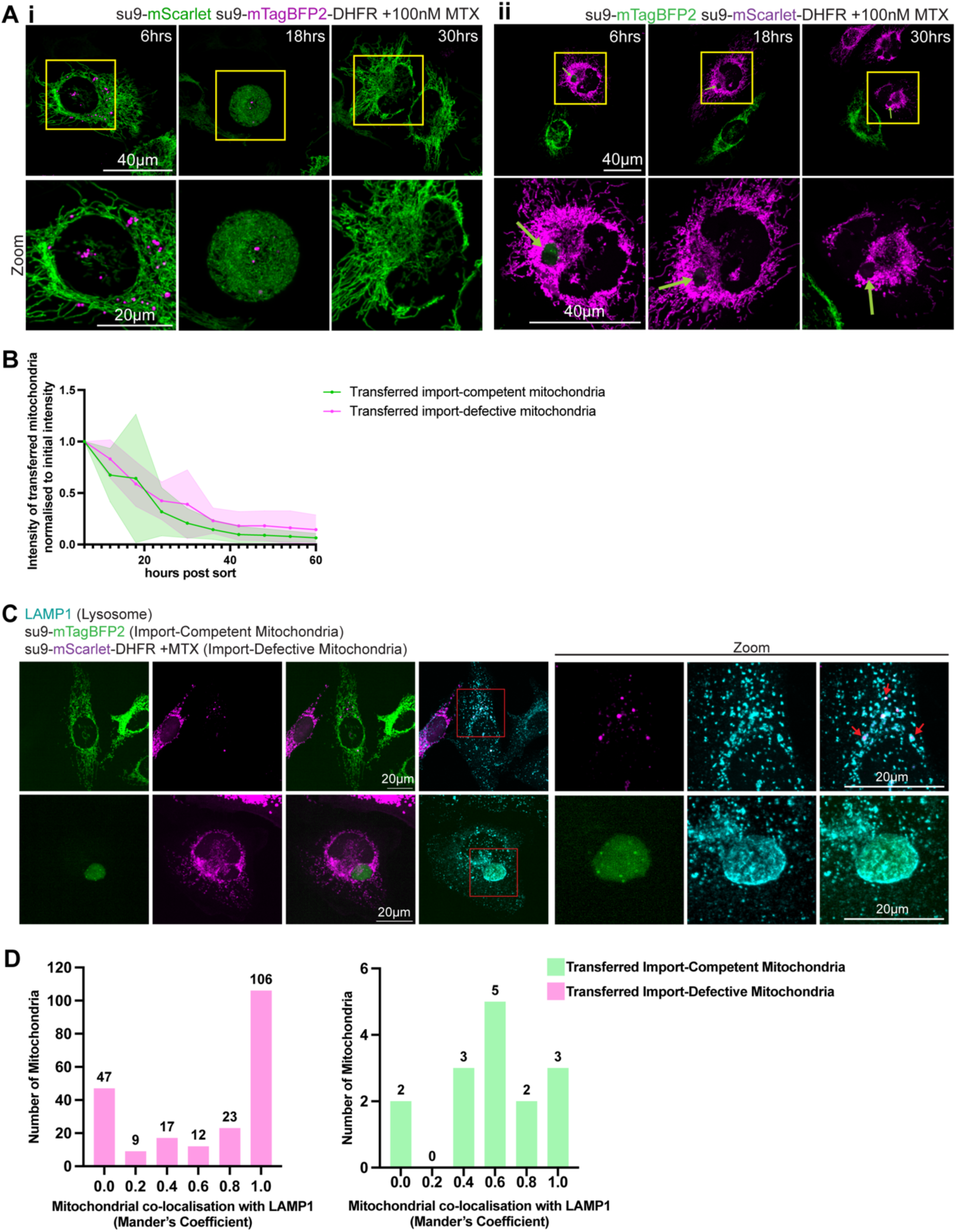
Transferred mitochondria are ultimately degraded. (Ai) Stills from a representative movie of transferred import-defective mitochondria (su9-mTagBFP2-DHFR +100nM MTX) in a cell containing import-competent mitochondria (su9-mScarlet) at indicated time points post-sorting. (Aii) Stills from a representative movie of transferred import-competent mitochondria (su9-mTagBFP2) in a cell containing import-defective mitochondria (su9-mScarlet-DHFR +100nM MTX) at indicated time points post-sorting. (B) Quantification of intensity of transferred import-defective mitochondria (su9-mTagBFP2-DHFR +100nM MTX) normalized to intensity at 6hrs post sorting (n=11 cells) and quantification of intensity of transferred import-competent mitochondria (su9-mTagBFP2) normalized to intensity at 6hrs post sorting (n=7 cells). Lines plotted from mean normalized intensity for each timepoint and area shading shows standard deviation. (C) Representative images of LAMP1 stained double positive cells from a 48hr coculture of su9-mTagBFP2 and su9-mScarlet-DHFR cell lines in the presence of MTX. (D) Histogram of Mander’s coefficient for mitochondrial co-localization with LAMP1. n=47 cells from N=3 biological replicates.

The fluorescence intensity of the transferred mitochondria over the 60hr imaging period was then calculated and plotted relative to the initial intensity (Figure 5B). Transferred import-defective mitochondria showed an intensity of 52.2% of the initial intensity at 18hrs post sorting, 19.5% at 36hrs, and 9.13% at 60hrs. Similarly, transferred import-competent mitochondria exhibited an intensity of 41.0% of the initial intensity at 18hrs post sorting, 11.6% at 36hrs, and 5.38% at 60hrs. These reductions in fluorescence intensity is indicative of lysosomal degradation. Co-localisation of transferred import-defective and import-competent mitochondria with the lysosomal marker LAMP1 confirmed this (representative images in Figure 5C and quantification in D). Overall, our results show that transferred mitochondria, regardless of their fragmentation and dispersal or accumulation within MDBs, are degraded by their recipients at a similar rates.

### Conclusions

We show here that bidirectional intercellular transfer of import-competent and import-defective mitochondria *via* TNTs ultimately leads to transmitophagy. We hypothesise that the different directions of traffic establish distinct, but complementary, mechanisms for the rescue of mitochondrial import failure (Figure 6). On the one hand, we suppose the intercellular trafficking of import-defective mitochondria, are simply for the purpose of transmitophagy (Davis *et al*, 2014). In these cells the autophagy capacity is likely overwhelmed, with export of compromised mitochondria serving to outsource the degradative burden to another cell, noted also elsewhere (Chakraborty *et al*, 2025). This process may be particularly important for cells prone to high levels of mitochondrial import dysfunction or with low mitophagy capability. On the other hand, we suggest that transfer of import-competent mitochondria is a response to extracellular stress signalling from cells containing import-defective mitochondria. Similarly to what has been reported before (Lin *et al*, 2024; Kidwell *et al*, 2023), the degradation of these mitochondria potentially functions to instigate additional events in the import-defective recipients to help facilitate their recovery by an as yet unknown mechanism. While further work will be required to test these possibilities, this work provides new insights of the role of canonical and non-canonical transmitophagy in mediating rescue of mitochondrial defects.

**Figure 6:**
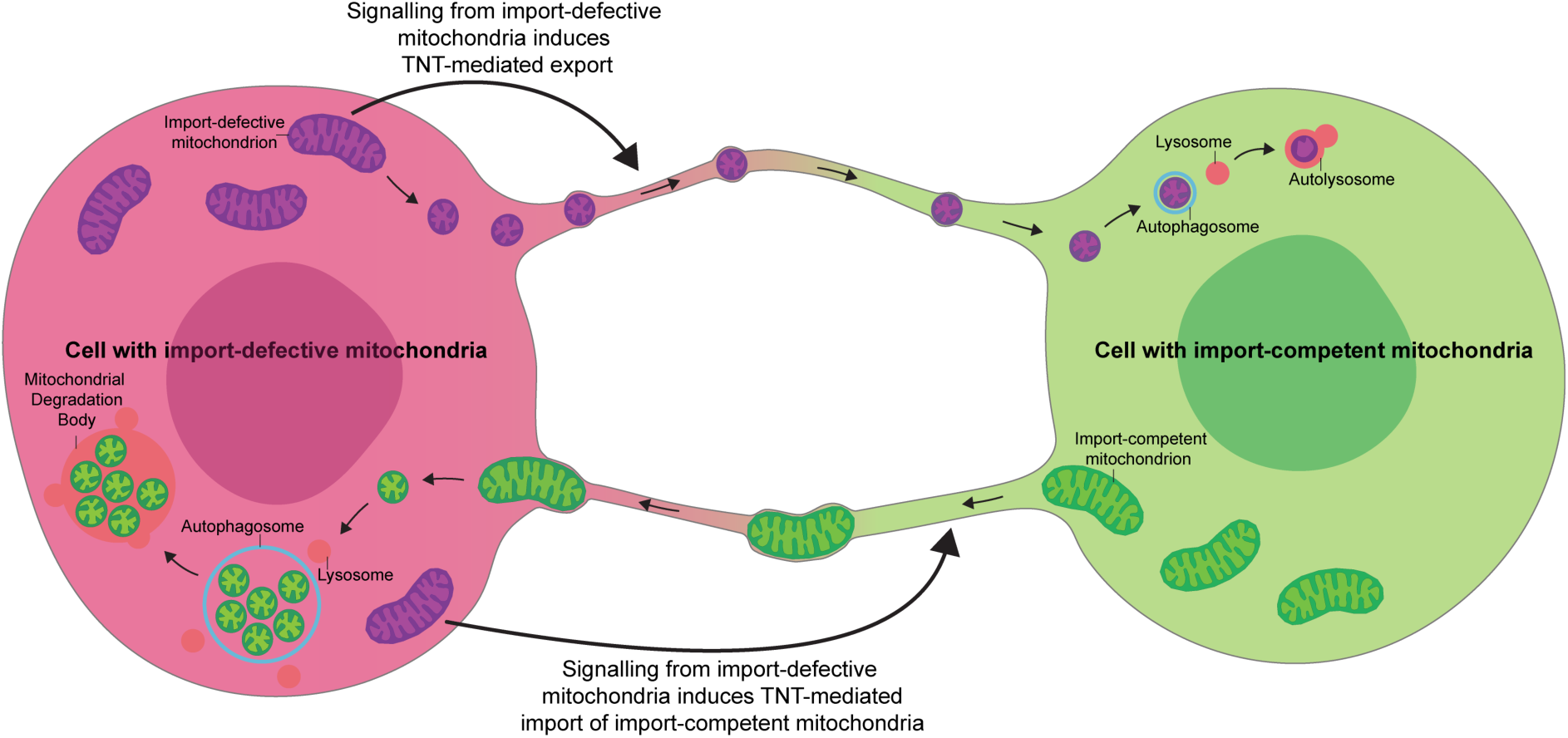
Schematic of model.

## MATERIALS AND METHODS

### Generation of constructs

Constructs were generated by standard cloning techniques and verified by Eurofins Genomics Whole Plasmid Sequencing service.

### HeLa cell culture

All cell lines were maintained in T75 ventilated flasks in humidified incubators at 37°C with 5% CO2. HeLa cells were ‘conditioned’ in galactose media (DMEM [+] L-Glutamine (gibco), 10mM D-Galactose, 1mM Sodium Pyruvate, 10% FBS) for three weeks, washing with 1 x PBS every 3 days. After this point, cells were considered OXPHOS dependent and termed ‘HeLaGAL’ cells (Aguer *et al*, 2011; Marroquin *et al*, 2007). Cells were split 1:5 every 2-3 days.

### Primary astrocyte cell culture

Primary astrocytes were isolated from embryonic day 17 (E17) Han Wistar rat pups. Pregnant Han Wistar rats (Charles River) were anaesthetised using isoflurane with pure oxygen flow and humanely killed by means of cervical dislocation, following Home Office Schedule 1 regulations. Isolated cortices were washed extensively in HBSS and dissociated by incubation with 10% (v/v) Trypsin-EDTA solution at 37°C for 15 mins.

Cortical cells were seeded at a density of 1 x 10^7^ cells in Poly-L-Lysine-coated T75 flasks. Astrocytes were grown in DMEM supplemented with 10% (v/v) fetal bovine serum (FBS), 1% P/S and 5 mM glucose maintained in humidified incubators at 37°C with 5% CO2. The media was replenished every 2-3 days and astrocytes were passaged every 7-8 days. On day 7, microglia were removed using an orbital shaker (180 rpm for 30 min) followed by the removal of oligodendrocytes (300 rpm for 6 hr). Primary astrocytes were maintained in culture for up to 4 weeks.

All animal care and procedures were carried out in full compliance with University of Bristol and ARRIVE guidelines, and the UK Animals Scientific Procedures Act, 1986. In addition, all experimental protocols were approved by University of Bristol Animal Welfare and Ethics Review Body (ethics approval number UIN: UB/23/069) panel and the Biological and Genetic Modification Safety Committee (BGMSC).

For mitochondrial transfer experiments astrocytes were infected with lentivirus encoding su9-EGFP-DHFR or Mito-DsRed. After 7–14 days of expression, co-cultures were established. Astrocytes were washed several times with PBS, dissociated with Trypsin-EDTA for 3 min at 37 °C, and the enzymatic activity quenched with complete media. Mito-DsRed– and su9-EGFP-DHFR–expressing astrocytes were mixed and plated onto 1× Geltrex-coated coverslips. Co-cultures were incubated for 24 hrs at 37 °C with 5% CO₂, followed by treatment with 100 nM MTX for 48 hrs to block mitochondrial protein import in su9-EGFP-DHFR–infected cells.

### Lentivirus production, transduction and selection

Lentiviral particles were produced in HEK293T cells using the construct of interest in pLVX vector, alongside packaging vectors pAX2 and pMDG2, in combination with pEI transfection reagent.

HEK293Ts were washed with 1 x PBS then the PEI/DNA solution was added and cells were incubated for 4 hours at 37°C 5%CO2. Following this, the PEI/DNA solution was removed and replaced with complete media.

Lentivirus was harvested after 72hrs. Media was removed and spun at 4000rpm for 5 minutes to pellet dead cells and the supernatant was then filtered through a 0.45um filter and added to HeLaGALs or aliquoted and stored in the −80°C for future use.

Infected HeLaGALs were incubated for 72hrs prior to selection. Cells were washed in 1 x PBS and complete media containing 3μg/mL puromycin was added. This was repeated every day until no cells were left in the control well. Remaining cells in the infected wells were grown up for experimental use.

### Fluorescence-activated cell sorting

Cells were washed in 1 x PBS then incubated with room temperature Accutase for 15 minutes. Detached cells were pelleted by centrifugation at 300xg for 2 minutes, resuspended in ice-cold MACS buffer (1 x PBS, 5mM EDTA, 0.5% BSA) then incubated with DRAQ7 Dye prior to cell sorting.

Cell sorting was performed on the BD FACS Aria II or the BD Influx sorters.

Parental HeLaGAL cells were used establish a gating strategy for the isolation of live singlet cells. Following this, mono-cultures of HeLaGAL cells expressing either su9-mScarlet-DHFR, su9-EGFP or su9-mTagBFP2 were used to set up a four-way gating system in which the live singlet cells containing both mScarlet or EGFP could be sorted.

Sorted cells were collected in 50%FBS + 50%PBS with 1% Penicillin-Streptomycin (Gibco), pelleted by centrifugation at 300xg for 2 minutes then resuspended in complete media with 1% Penicillin-Streptomycin (Gibco).

### Florescence microscopy

All florescence microscopy of cells was performed on an Olympus SpinSR spinning disk microscope.

### LAMP1 staining

Cells were fixed in 4% PFA for 10 minutes then washed 3 times in 1 x PBS prior to permeabilization with 0.1% saponin for 5 minutes at room temperature. Cells were blocked in 5% BSA in PBS for 1hr then incubated with LAMP1 antibody (H4A3, DSHB) in 5% BSA in PBS for 1 hour. Coverslips were then washed 3 times for 5 minutes in 1 x PBS then incubated with secondary antibody in 5% BSA in PBS for 1 hour. Coverslips were then washed a further 3 times for 5mins in 1 x PBS and a final wash in H2O prior to mounting on slides with Fluoromount-G (Invitrogen).

### DAPI staining

Cells were fixed in 4% PFA for 10 minutes then washed 3 times in 1 x PBS and a final wash in H2O prior to mounting on slides with with Fluoromount-G with DAPI (Invitrogen).

### Live Imaging Probes

1000x stock of CellMask (Invitrogen) was diluted in complete media. Cells were incubated with this for 10 minutes, washed in 1 x PBS then put into imaging media.

### Correlative light and electron microscopy

Sorted cells were seeded onto 35mm dishes with a no. 1.5 gridded coverslip, 14mm glass diameter (Mattek) in complete media, 100nM MTX, 1% Penicillin-Streptomycin and left overnight to adhere. Cells were live imaged on a Spinning disc microscope and fluorescence images taken along with a DIC overview of grid coordinates.

Samples were then fixed in 2.5% glutaraldehyde in 0.1M cacodylate buffer (pH 7.3) for 30 minutes at room temperature. Dishes were washed in 0.1M cacodylate buffer before post fixing in 1% Osmium, 1% Potassium Ferrocyanide in 0.1M cacodylate buffer for 1 hour at room temperature. Dishes were then washed in water and stained with 3% uranyl acetate for 1 hour at room temperature prior to rinsing 3 x in water and then dehydrated through a graded ethanol series (50%, 70%, 80%, 90%, 96%, 100%, 100%) for 5 minutes each. Samples were then infiltrated in Epoxy resin x 3 (1 hour each) before polymerisation at 60°C for 48 hours.

Coverslips were removed by immersing dishes in liquid nitrogen followed by boiling water. Samples were trimmed to ROI with razor blades. 70nm (thin) and 250nm (thick) sections were cut on an Ultramicrotome (UC7, Leica) using a diamond knife (Diatome) and mounted onto formvar coated slot grids. Thick sections were incubated in 15nm gold fiducials (Aurion) for 5 minutes on each side before imaging for tomography.

Images were acquired at 120kV on a Talos L120C transmission electron microscope equipped with Lab6 filament and 4K x 4K Ceta camera. Tomograms were reconstructed and combined using IMOD software [3,4].

Fluorescence images were correlated with electron microscopy images using easy cell-correlative light to electron microscopy (eC-CLEM) (Paul-Gilloteaux *et al*, 2017) in the Icy platform (de Chaumont *et al*, 2012).

### Light microscopy image processing and analysis

Images were processed and analysed on Fiji ImageJ (Schneider *et al*, 2012) using ModularImageAnalysis (MIA) (Cross *et al*, 2024). GraphPad Prism software was used for all statistical analyses and graph design.

## ACKNOWLEDGEMENTS

The authors gratefully acknowledge Dominic Alibhai of the Wolfson Bioimaging Facility for their assistance in this work.

## AUTHOR CONTRIBUTION

EG and IC devised the project. EG designed experiments. EG and BW produced and analysed the data. EG interpreted the data. EG, KW and IC wrote the manuscript. IC secured funding and led the project.

## DECLARATION

The authors declare no competing interests.

## DATA AND MATERIALS AVAILABILITY

All data are available in the main text.

